# An *In-Silico* Mammalian Whole-Cell Model Reveals the Influence of Spatial Organization on RNA Splicing Efficiency

**DOI:** 10.1101/435628

**Authors:** Zhaleh Ghaemia, Joseph R. Peterson, Martin Gruebele, Zaida Luthey-Schulten

**Affiliations:** Department of Chemistry, University of Illinois at Urbana-Champaign, Urbana, IL, 61801; Center for the Physics of the Living Cells, University of Illinois at Urbana-Champaign, Urbana, IL, 61801; Beckman Institute, University of Illinois at Urbana-Champaign, Urbana, IL, 61801.

**Keywords:** Human whole-cell modeling, RNA splicing, Stochastic reaction-diffusion simulations

## Abstract

Spatial organization is a characteristic of eukaryotic cells, achieved by utilizing both membrane-bound and non-bound organelles. We model the effects of this organization and of organelle heterogeneity on RNA splicing (the process of making translationally-ready messenger RNA) and on splicing particles (the building blocks of splicing machinery) in mammalian cells. We constructed a spatially-resolved whole HeLa cell model from various experimental data and developed reaction networks to describe the RNA splicing processes. We incorporated these networks into our whole-cell model and performed stochastic simulations for up to 15 minutes of biological time. We find that the number of nuclear pore complexes affects the number of assembled splicing particles; that a slight increase of splicing particle localization in nuclear speckles (non-membrane-bound organelles) leads to disproportionate enhancement in the mRNA splicing and reduction in the transcript noise; and that compartmentalization is critical for a correctly-assembled particle yield. Our model also predicts that the distance between genes and speckles has a considerable effect on effective mRNA production rate, further emphasizing the importance of genome organization around speckles. The HeLa cell model, including organelles and subcompartments, provides an adaptable foundation to study other cellular processes which are strongly modulated by spatio-temporal heterogeneity.

**Significance Statement:** The spliceosome is one of the most complex cellular machineries that cuts and splices the RNA code in eukaryotic cells. It dynamically assembles, disassembles, and its components are formed in multiple compartments. The efficiency of splicing process depends on localization of its components in nuclear membrane-less organelles. Therefore, a computational model of spliceosomal function must contain a spatial model of the entire cell. However, building such a model is a challenging task, mainly due to the lack of homogeneous experimental data and a suitable computational framework. Here, we overcome these challenges and present a whole HeLa cell model, with nuclear, subnuclear, and extensive cytoplasmic structures. The three-dimensional model is supplemented by reaction-diffusion processes to shed light on the function of the spliceosome.

---

Cells use spatial organization to mediate the complex biochemical reaction networks. Although membranes have long been recognized as means to confine organelle-specific compounds, non-membrane-bound organelles are increasingly found to play crucial roles in cellular functions (1). These organelles can be formed as liquid-liquid phase separated regions and are therefore also known as liquid droplets (2). Cells may have numerous such liquid droplets that form either in the cytoplasm or the nucleus (3, 4). Each droplet is involved in specific cellular processes. As a prime example, nuclear speckles, or interchromatin granules, are droplets formed in the nucleus that are thought to be primarily involved in pre-mRNA splicing (5).

RNA splicing has evolved in eukaryotic cells to allow cell complexity without massively increasing gene count. Instead, the structure of genes changed such that the coding regions (exons) are interrupted by non-coding regions (introns) (6). Coding regions must then be ligated, to form functional transcripts. There are, on average, eight introns per gene (7), so coding regions can be shuffled after the removal of all or a subset of introns by a process called alternative splicing. The order of intronic removal defines the function of the protein coded by the transcript; thus, a single gene can encode a variety of functionalities (8). Spliceosome is the cellular machinery that binds to the intron/exon sites, removes the introns and joins the exon ends. It is a multi-megadalton complex consisting of five (uridine rich) protein-RNA small nuclear ribonucleoprotein (snRNP) complexes: U1 snRNP, U2 snRNP, U4 snRNP, U5 snRNP and U6snRNP (9). The biogenesis of splicing particles (snRNP complexes) occurs in multiple steps in both nucleus and cytoplasm, and finalizes in the nucleus (6). The mature splicing particles then localize in nuclear speckles and assemble on the pre-messenger RNA (pre-mRNA) transcripts in a coordinated and step-wise fashion, and upon completion of the splicing reaction, they disassemble (10).

Klingauf et al. showed that the association of splicing particles, U4snRNP and U6snRNP in Cajal bodies (another type of phase-separated nuclear regions) is enhanced compared to the assembly reactions taking place throughout the nucleus, pointing to the importance of cellular organization even at the sub-nuclear level (11). Chang and Marshall proposed in a commentary that organelle heterogeneity can also lead to cellular phenotypic behavior, similar to the heterogeneity at the molecular level (12). Additionally, a very active effort is underway to determine the cellular organization, such as the one carried out by Johnson et al. on human induced pluripotent stem cells, from a rich set of cellular fluorescence images (13).

Although the basic utility of nuclear speckles in pre-mRNA splicing is already appreciated, the influence of spatial localization on splicing activity and mRNA production is not understood at a quantitative level. The effect of variations in the involved organelles has not been investigated either. Here, we construct a 18-*μm* spatially-resolved model of a whole mammalian cell, specifically a HeLa cell, from a library of experimental data such as cryo-electron tomography (14), mass spectrometry (15), fluorescent microscopy and live-cell imaging (11, 16–18) and -omics data (19, 20). We simulate our eukaryotic cell with organelles, compartments and biomacromolecules, within the framework of reaction-diffusion master equations with Lattice Microbes software (21, 22) for up to 15 minutes of biological time. Our simulations explore how cellular organization affects the efficiency of spliceosomal particle formation and pre-mRNA splicing. Specifically, we find that even a slight increase in the relative localization of splicing particles in nuclear speckles can both enhance mRNA production and reduce its noise. Additionally, we rationalize the biological selection of design parameters of nuclear speckles, specifically, their size and number. Finally, we predict that the organization of active genes around nuclear speckles can affect mRNA production.

## Results and discussion

### Spatially-resolved model of a HeLa cell

We used a data-driven approach to construct a representative HeLa whole-cell model that has not been available thus far. First, we gathered structural and -omics information from a variety of experimental studies (11, 14–20, 23–26). Then, the assembled data was ensured to be consistent with protein composition percentages of HeLa cell organelles determined by mass spectrometry (15) (see Methods for details). On average, proteins are composed of similar C/N/O/H ratios, hence mass percentages of organelles are approximately similar to volume percentages.

Figure 1-A and B, show the overall HeLa cell model and a more detailed view of the nuclear region. Assuming experimental growth conditions resulting in spherically-shaped cells (25, 27), a volume of 3000 *μm*^3^ (23) leads to a cellular radius of 8.9 *μ*m. The essential components of the cell include: plasma membrane, cytoplasm, endoplasmic reticulum (ER), mitochondria, Golgi apparatus and nucleus. The ER units were modeled by stochastic shapes using a cellular automata algorithm (see Supplementary Information for details and algorithm). The ER units are distributed in the cytoplasm, spanning from the nuclear envelope to the plasma membrane, and are intertwined with other cytoplasmic organelles (28, 29). The ER units make up ∼ 4.5% of the cell volume (15). About 2000 rod-shaped mitochondria with dimensions of 0.6 *μ*m × 0.49 *μ*m were randomly placed throughout the cytoplasm, filling ∼ 11% of the total volume (15). A Golgi apparatus consisting of five stacked sheets, each with a thickness of 0.128 *μ*m, was placed close to the nucleus (30).

**Fig. 1.**
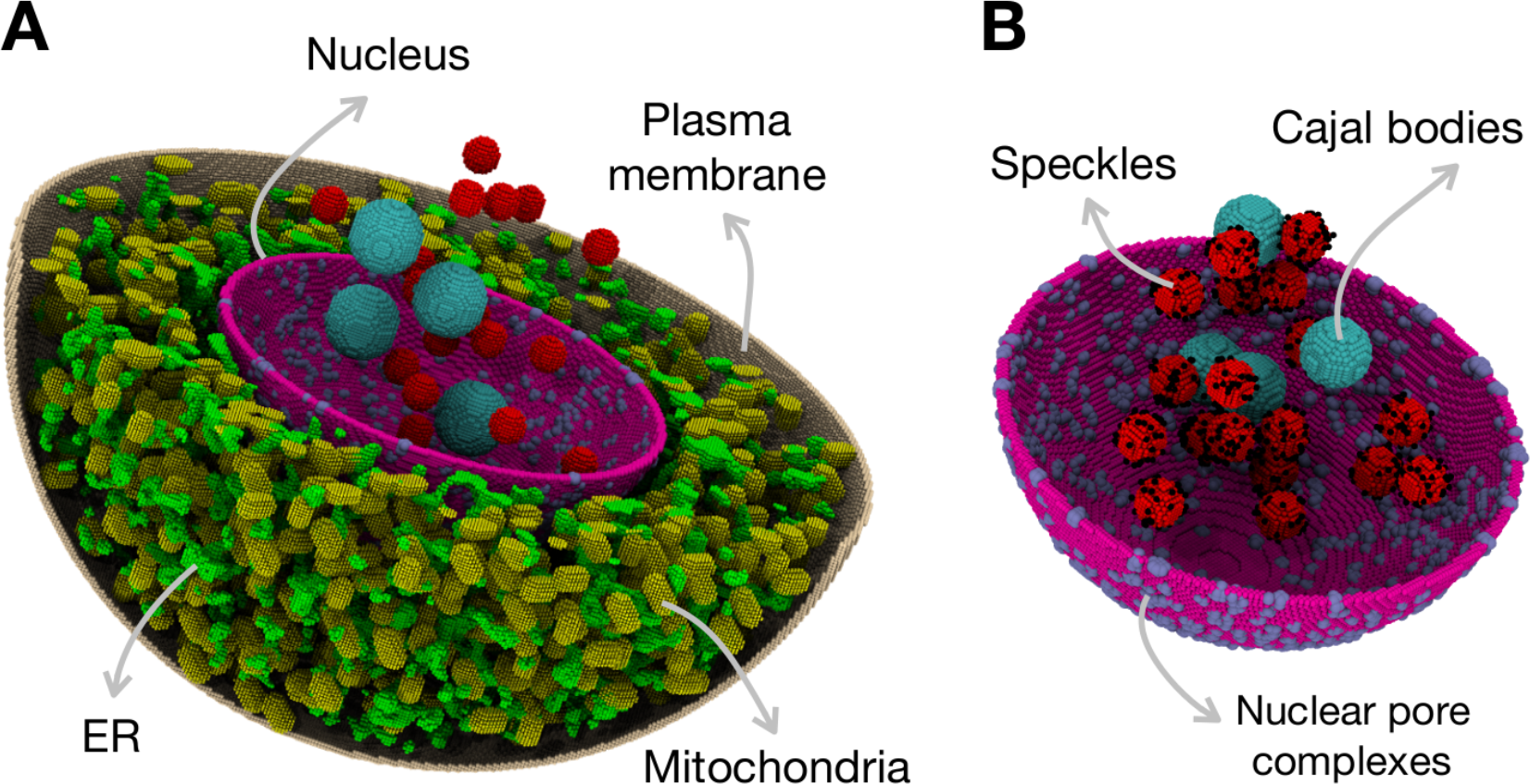
A data-driven model for a 18 *μm* HeLa cell: A) cytoplasmic components are: ER, mitochondria and Golgi (not shown); and B) nucleus containing nuclear pore complexes, Cajal bodies and nuclear speckles.

The nucleus, which plays a critical role in our model, has a radius in the range of 3.74–5.29 *μ*m (24–26). It consists of nuclear pore complexes (NPCs) of 0.083 *μ*m radii (14) and a density of 7 per *μm*^2^ (16), 20 spherically-shaped nuclear speckles with 0.35 *μ*m radii (18) and 4 Cajal bodies with 0.5 *μ*m radii (11). Active genes (black dots in Figure 1-B) were placed around the speckles (31–33). The nuclear components were chosen mainly among those that play a role in RNA splicing processes. Cellular components included in the *in-silico* model are listed in Table 1 along with their dimensions. The details of the construction of each organelle are provided in the Methods section.

**Table 1.** The cellular components of the constructed HeLa model.

| Component | Dimension ($\mu\text{m}$) | Number | Reference |
| --- | --- | --- | --- |
| HeLa cell | $R = 8.9$ | 1 | (23) |
| Nucleus | $R = 3.74, 4.15, 4.67, 5.29$ | 1 | (15, 24–26) |
| Nuclear pore complexes | $R = 0.083$ | 1230, 1515, 1918, 2461 | (14, 16) |
| Mitochondria | $0.6 \times 0.49$ | 2000 | (17) |
| Nuclear speckles | $R = 0.35$ | 20 | (18) |
| Cajal bodies | $R = 0.5$ | 4 | (11) |
| ER | – | 4.5% cell volume | (15) |

### The kinetic model for spliceosome formation and action

We studied two processes: first, the formation of splicing particles (U1snRNP and U2snRNP) which is a multi-compartmental process and second, the spliceosome assembly, splicing reaction and generation of mRNA transcripts. Together, these capture the whole process of splicing from machinary construction to functional transcript production. After the assembly of U1snRNP and U2snRNP in our model (the first process), the pre-mRNA transcripts are spliced (the second process) according to the following reduced scheme for the spliceosome assembly:

1. An active-28 Kb gene is transcribed and pre-mRNA transcripts are produced
2. U1snRNP and U2snRNP particles are formed and are present in the cell nucleus
3. U4/U6.U5 trisnRNP particles are also present in the nucleus; Because of the complexity in the formation of these complexes we assumed they are pre-formed in our model (34).
4. The spliceosome assembles in a stepwise manner on pre-mRNA transcripts
5. After splicing occurs, the pre-mRNA is converted to an mRNA transcript
6. The spliceosome disassembles after splicing, ready to assemble on another transcript

Below, we describe in details the splicing assembly and reaction and splicing particles formation.

#### Formation of splicing particles

A splicing particle consists of a uridine-rich small nuclear RNA (U snRNA) that is bound to a heptamer ring of proteins, called Smith proteins (Sm) and variable numbers of particle-specific proteins. The formation of splicing particles happens in multiple steps and compartments. To understand the effects of geometry on the formation process, we developed a kinetic model to describe these processes and studied them in our developed spatially-resolved HeLa cell model.

Figure 2 shows the steps associated with the formation of splicing particles and the reactions are summarized in Table 2 in Methods. Upon transcription, U1(2) snRNA has to pass through nuclear pore complexes to reach cytoplasm, where by a series of complex reactions they bind to Sm proteins. Inspired by two studies (35, 36), we proposed the following mechanisms for the cytoplasmic part of the process: U1(2) snRNA transcript binds to Gemin 5 (*G*^5^) which is part of the survival of motor neurons (SMN)-Gemin complex that mediates the Sm proteins assembly on snRNA. The formed complex then binds to a ring of five already-assembled Sm proteins (*Sm*^5^) through a process called RNP exchange suggested by Ref. (36). This process facilitates the Sm proteins binding to the snRNA transcript and the release of *G*^5^. In the last step, the remaining Sm proteins (*Sm*^2^) joins the complex and the *U*1(2)*snRNA.Sm*^7^ complex is formed. After the completion of the binding of Sm proteins on snRNA, the complex again pass through the NPCs and make its way to the nucleus. At nucleus, the 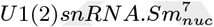 complex localizes to the Cajal bodies (37) and binds to particle-specific proteins; *U*1(2)_*prot*_, and the mature splicing particle is formed.

**Fig. 2.**
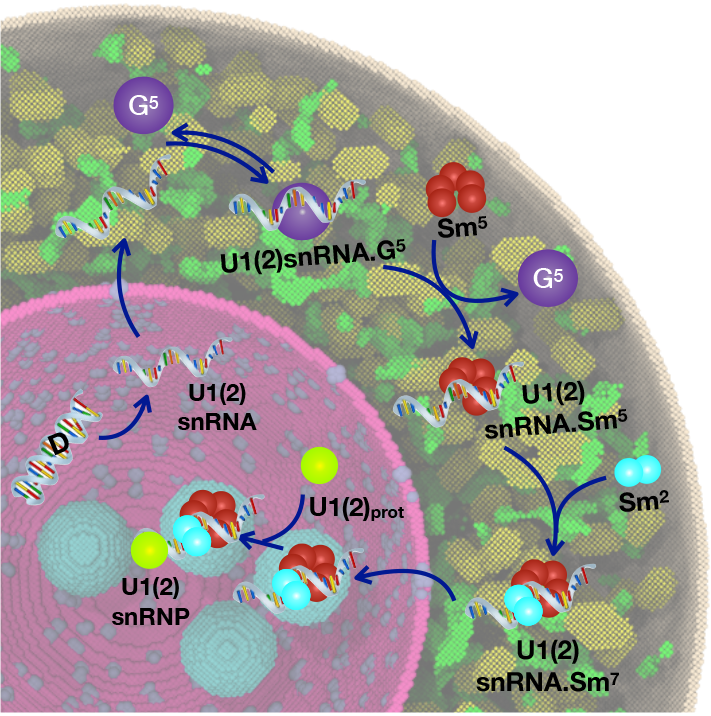
Reaction scheme for describing the formation of U1 and U2 splicing particles mapped on a cross-section of our *in-silico* HeLa cell (35, 36). The four spherically-shaped regions in cyan color are Cajal bodies.

The diffusion coefficients for the species that are involved in the splicing particle formation reactions were mainly adopted from various experimental sources and are listed in Table S2.

#### Assembly of the spliceosome and splicing reaction

The assembly process of the spliceosome machinery is entangled with a complex network of auxiliary and regulatory proteins that detect the splice site and alternate the splice sites according to cellular cues by a process called alternative splicing (8). To simplify this network, we assume that a particular splice site has been chosen and focus only on the assembly of the spliceosomal particles on that site and the splicing reaction.

Figure 3 depicts our model for splicing reaction; and the details of the reactions and their associated rates are represented in Table 3 in Methods. According to the conventional splicoeosome assembly model (38), the U1snRNP particle binds to the 5′ end of the exon (“complex E”), following by binding of the U2 particle to the associate 3′ end to form “complex A”. To make a more realistic model, we added an additional initial reaction as suggested by Ref. (39): the U2snRNP can bind the pre-mRNA before U1snRNP, making “complex E*”. Regardless of the binding order of these splicing particles, a viable complex A is formed that can continue the remaining assembly process. The “tri.U” (U4/U6 bound to U5) then joins the complex forming “complex B”. Subsequently, U1snRNP leaves the complex for the catalytically active “complex B*”. The intron is then removed and the splicing particles are recycled for another round of assembly.

**Fig. 3.**
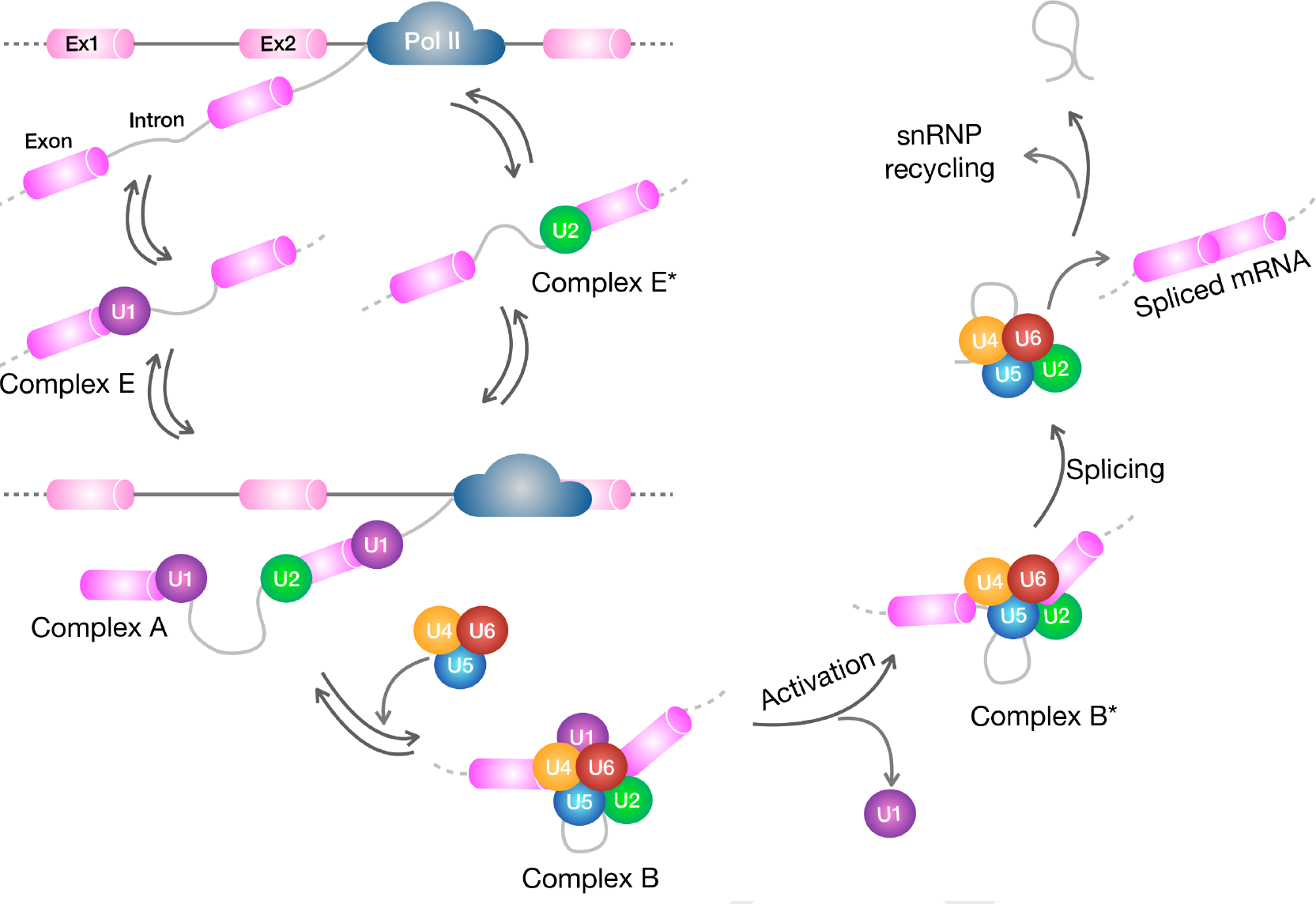
Splicing reactions as implemented in our simulations. The reactions together with their corresponding rate constants are shown in Table 3. Abbreviations are: Pol II (RNA polymerase II), Ex1 and 2 (Exon 1 and 2).

##### Co-transcriptional splicing

Splicing is known to be overwhelmingly co-transcriptional; meaning, as transcription is occurring, the spliceosome assembles on the transcribed pre-mRNA and splicing reactions begin. In our model, an average gene consisting of 8 introns and an intron length of 3.4 Kbase (plus 137 base for each exon) is considered (7). After the transcription of the first exon-intron-exon piece, the splicing reaction starts as discussed above. Simultaneously, another intron-exon pair is transcribed, continuing the spliced transcript. The cycle repeats till the end of the gene.

The diffusion coefficients of spliceosomal particles are also listed in Table S3.

### Organelle heterogeneity influence the formation of splicing particles

Cellular phenotypic behavior arising from organelle heterogeneity is a subject worthy of study (12). We investigated how heterogeneities in NPC count and nuclear size affect the formation of splicing particles. All splicing particles, except U6, are complexes of uridine-rich small nuclear RNA bound to a heptameric ring of Smith (Sm) proteins, along with specific proteins that bind to each splicing complex. Among the five particles which are required for spliceosome function, we focus on the first two (U1snRNP and U2snRNP) that start the spliceosome assembly. As shown in Figure 2 and described above, these particles are formed in a multi-compartmental process (6). Because, the components of splicing particles have to assemble in both nucleus and cytoplasm, therefore, translocation through the NPCs is a critical step. Live cell imaging showed that NPC count varies (by 10%) (16), and so does nuclear size (15, 24–26). We posited that these variations could influence the formation of splicing particles, which we tested by varying NPC count and nuclear size, and examining the effect on the number of particles formed after 30 seconds of biological time. Figure 4 shows that increasing (decreasing) the number of NPCs by 20% results in an increase (decrease) in the number of mature U1 and U2 splicing particles. This effect is consistent across the tested nuclear radius with range of 3.74–5.29 *μ*m (15, 24–26). It is found that the number of splicing particles formed does not change significantly with nuclear size. This can be explained by the fact that a larger nucleus has a larger number of NPCs, since the density of NPCs is constant. Consequently, longer diffusion times in a larger nucleus are compensated by shorter translocation times required when there are more NPCs.

**Fig. 4.**
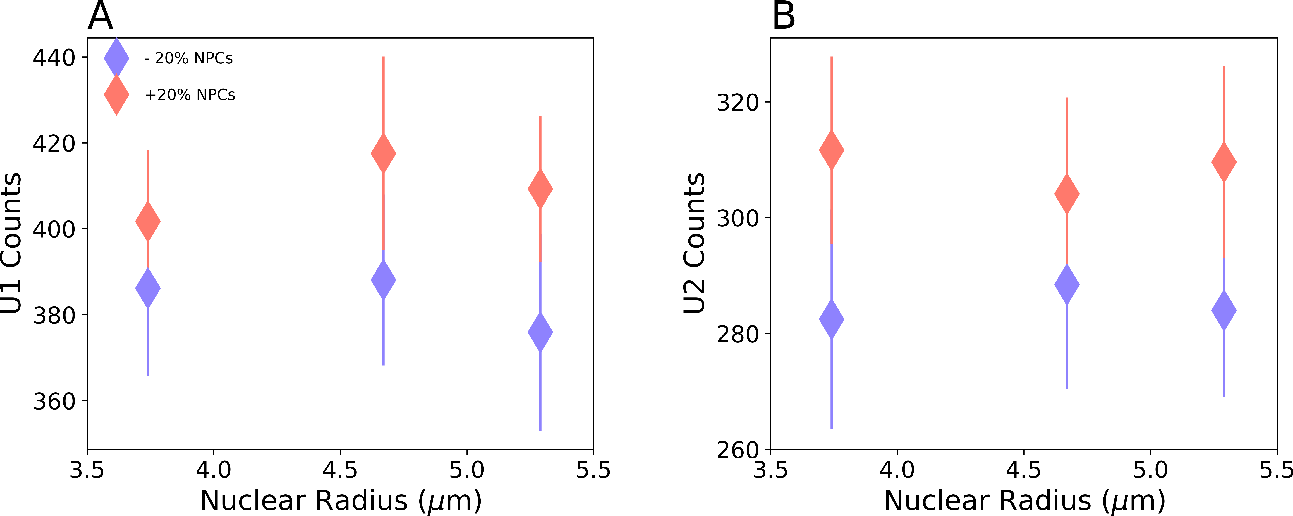
Spliceosomal particle formation depends on NPC count. Increase (pink) or decrease (blue) in the number NPCs by 20% results in a corresponding change in the number of U1 (A) and U2 (B) particles formed. The effect is consistent for different nuclear sizes. P-values for U1 results are: 1.4 × 10^−2^ (3.74 *μ*m), 1 × 10^−4^ (4.67 *μ*m), 9.6 *×* 10^−4^ (5.29 *μ*m); and for U2 results are: 9.5 × 10^−6^ (3.74 *μ*m), 8.3 × 10^−3^ (4.67 *μ*m), 1.3 × 10^−5^ (5.29 *μ*m). The cellular geometry in these simulations is the same as described in Table 1, except for the ER volume, which is ∼ 7% of the cell volume. However, the slightly higher occupancy is not known to have a considerable effect. Error bars represent the standard deviations. For each condition, 20 simulation replicates were performed.

To obtain insight into the formation of splicing particles, we dissected the overall kinetics of the process in terms of discrete reactions occurring in each compartment, i.e., nucleus and cytoplasm. These reactions include the transcription of snRNA and formation of (*U*1*snRN A*_*nuc*_), cytoplasmic production of U1snRNA Sm^7^, and finally the assembly of mature U1snRNP. We determined the timescale for the formation of each of these three species within the first assembled U1 particle. As shown in Figure 5, the series of cytoplasmic reactions take the longest to complete, irrespective of nuclear size. The timescale for cytoplasmic reactions is also statistically similar regardless of whether the ER or mitochondria or both are absent: for a full cell the time is 0.68 ± 0.30 s, as compared to 0.63 ± 0.35 s, 0.69 ± 0.24 s and 0.59 ± 0.21 s, respectively for cases where the cell model lacks ER, mitochondria, or both.

**Fig. 5.**
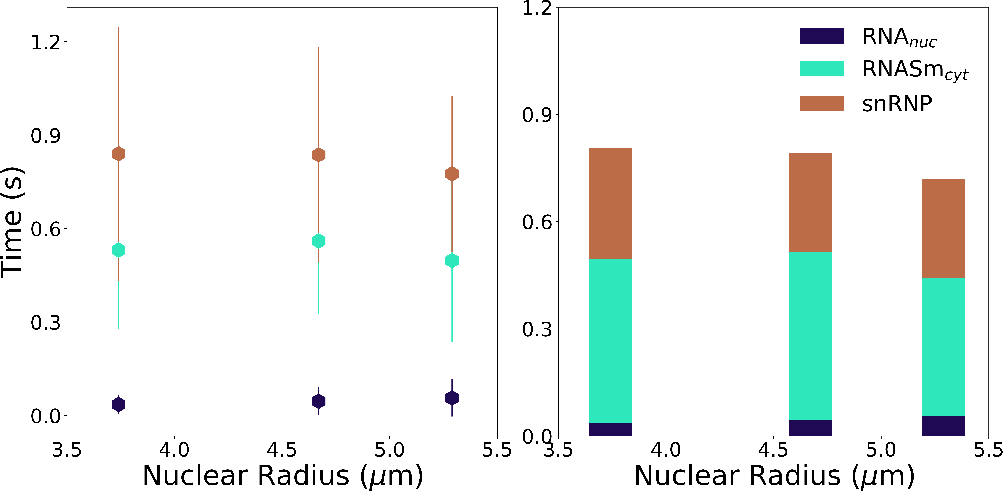
The time required to form the first splicing particle dissected by the set of discrete reactions occurring in the nucleus and cytoplasm. Error bars represent the standard deviations. For each condition, 20 simulation replicates were performed.

As mentioned above, in higher eukaryotes, different components of the splicing particles join the assembly in different compartments (6). This separation likely allows for higher quality control and prevents mixing of the partially-assembled particles with their substrates, thus preventing partially formed spliceosomes from deleteriously modifying pre-mRNAs. We examined the importance of multi-compartmentality by allowing all particle assembly steps to occur solely in the nucleus. We postulated that the latter modification may result in snRNA binding to proteins in an incorrect order, or, in incomplete assembly of the particle. In addition to confining assembly steps to the nucleus, we also modified the reaction network and added an extra reaction (*U*1*snRN A ․ Sm*^5^ + *U*1_*prot*_ → *U*1*RNA · Sm*^5^) to the set of reactions shown in Table 2, to account for misassembled splicing particles. The assembly of Sm-core is followed by RNA modification that triggers the nuclear import of the snRNA bound to Sm core (6). Therefore, in the multi-compartmental assembly process of splicing particles the U1snRNA · Sm^5^ complex is not found in the nucleus. As an outcome of simulating the nuclear assembly of the splicing particles, we found that although the system can make fully assembled splicing particles, it produces significantly more misassembled particles (771 ± 25) as compared to mature particles (248 ± 15), since the former are not required to go through the full assembly cycle (see Figure 2). This simulation result demonstrates the critical need for the compartmentalization of the overall assembly of the splicing particles. Similar multi-compartmental processes have been observed for other cellular machines such as ribosomal subunits (6).

### Nuclear speckles enhance effective splicing rate and control noise in mRNA

Nuclear speckles are self-organized liquid droplets that are known to act as stores for splicing particles in addition to other processes such as DNA repair and RNA modifications (5, 18). There is evidence that a subset of splicing occurs within or at the periphery of these nuclear bodies (40). Nuclear speckles, being liquid-liquid phase-separated regions, promote certain biochemical reactions, which is suggested to be due to an enhanced concentration of the reactants (2). To examine this phenomenon, we developed a reaction network to account for spliceosome assembly on pre-mRNA transcripts and splicing reaction described in details previously (see Figure 3). This network was included in stochastic simulations containing speckles in the HeLa cell and we determined the resulting effect on mRNA production and noise. The speckles were comprised of a concentrated store of splicing particles produced in the following manner (see Methods section). Briefly, we set the probability, P_n_, for splicing particles to transit from the nucleus into the speckles higher than the probability, P_s_, for the reverse transition. We found that the higher this imbalance (P_n_/P_s_) is, the greater the degree of localization of splicing particles in the speckles (see Figure 6-A). We compared mRNA production in cells with different degree of splicing particle localization in the speckles and also that in a control cell containing no speckles but with splicing particles randomly distributed throughout the nucleus. Relative to the case of no speckles (U1 fraction = 0 in Figure 6), a cell with about 10% of U1 located in speckles showed a large enhancement in the number of spliced mRNA transcripts from 0.25 to 40, which is effectively a ≈ 150-fold amplification (Figure 6-B,C). Thus, even a slight increase in the localization of splicing particles enhances mRNA production. The mRNA production continues to grow with further increase in the splicing particle localization up to an enrichment level of ≈ 55% U1 in speckles. Beyond this point, little increase in average mRNA count is observed (Figure 6-C). Alongside these trends, we examined the effect of speckles on the noise associated with mRNA production, estimated in terms of the coefficient of variation, *η*, which is the ratio of average mRNA counts to the standard deviation. As the percentage of splicing particles in speckles increases (Figure 6-D), the noise decreases. Thus, nuclear speckles not only enhance splicing activity, but they also help limit the noise that splicing introduces into the whole gene expression process. Using green fluorescent protein labels, Rino et al. determined the ratio of splicing protein U2AF in speckles to that in the nucleus to be 1.27 ± 0.07 (41). Strikingly, this experimentally determined ratio corresponds in our model to a *P*_*n*_/*P*_*s*_ = 100 (Figure 6-A), which is effectively a ∼ 55% localization of splicing particles in speckles, the point at which the mRNA production has maximized.

**Fig. 6.**
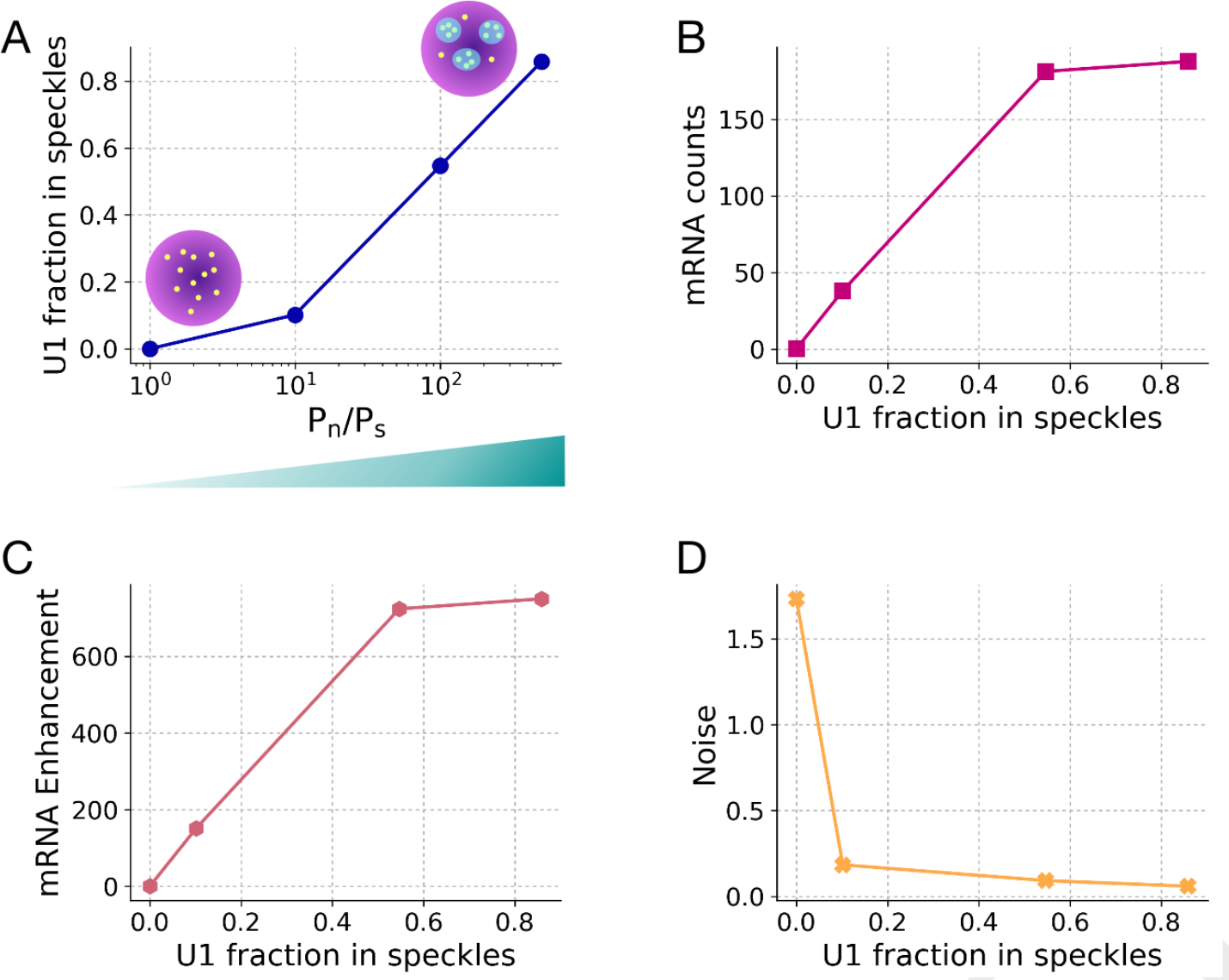
Splicing efficiency increases in the presence of speckles in the cells: A) The higher the probability for the splicing particles to transition from the cell nucleus to the speckles, relative to the reverse transition, the higher is the localization of splicing particles in speckles. Schematically, the randomly distributed splicing particles (yellow dots) in the cell nucleus (colored in purple), localize in nucleus speckles (blue shaded regions) as the probability imbalance increases. B) As the percentage of splicing particles located in speckles increases, the number of spliced mRNA also increases. C) This enhancement in mRNA production is highly sensitive to the localization of splicing particles in speckles: with only a 10% localization of splicing particles in speckles, the splicing reaction is enhanced ∼ 150-fold relative to the case with no speckles. D) Noise (average of mRNA counts/the standard deviation) decreases as a greater percentage of splicing particles are localized in speckles. For each condition, 20 simulation replicates were performed.

#### Speckle-enhanced splicing is concentration-dependent

The number of splicing particles required per pre-mRNA transcript is a function of many variables including the rate of transcription (7) and therefore this number may vary from one gene to another. We investigated how variation in the ratio of splicing particles to pre-mRNA transcripts affects overall mRNA transcript production and noise in a cell with nuclear speckles. Specifically, for 20 constitutively transcribing genes, we changed the number of particles available for pre-mRNA binding and splicing from 16 to 1600 corresponding to a concentration range of 0.1–10 nM. The remainder of the total 105 splicing particles (42) were bound to pre-mRNA transcripts and actively splicing. Figure 7-A summarizes how the concentration of U1 splicing particles affects the ability of speckles to enhance splicing. At 1 nM U1, mRNA production in a cell with speckles (with P_n_/P_s_ = 500) is ≈ 750-fold that for a cell with no speckles. At 10 nM, this enhancement factor reduces to 1.4-fold. Thus, at lower concentrations of U1, speckles enhance splicing much more strongly. Consistently, as figure 7-B shows, the noise of mRNA production is also influenced. At 1 nM U1, the noise in the presence of speckles is ≈ 30-fold lower as compared to a cell with no speckles; whereas, at 10 nM, the noise is unaffected by whether speckles are present or not.

**Fig. 7.**
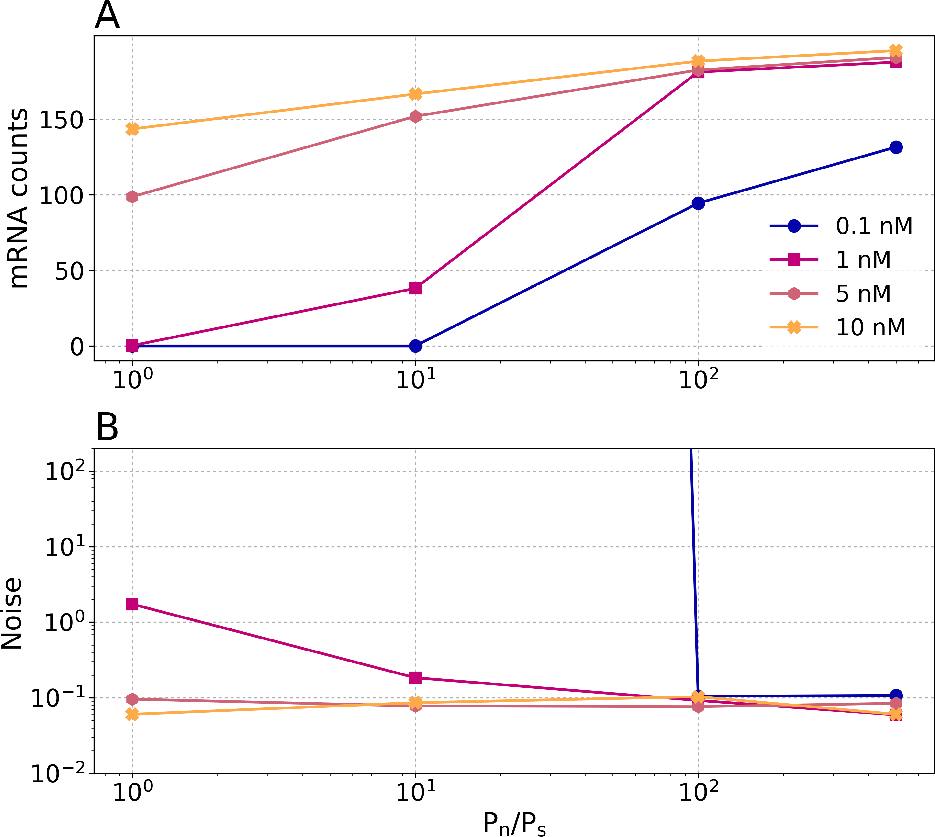
Splicing particle concentration affects the functional advantage of speckles: A) Enhancement in mRNA production due to the presence of speckles, depends on the U1 splicing particle concentration. B) Effect of the U1 splicing particle concentration on the mRNA production noise. For each condition, 20 simulation replicates were performed.

### Speckle size and number have been fine-tuned by cells to optimize mRNA production

We investigated how a cell decides the number and the size of nuclear speckles, after the dedication of certain percentage of its nuclear volume to these organelles. We hypothesize that the experimentally observed anatomy of the speckles is optimized by the cells. To test this hypothesis, we assigned about 10% of the nuclear volume to speckle (20) and keeping the total volume of speckles constant, we increased the number of speckles and reduced their sizes, as shown in Figure 8-A. Increasing the number of speckles results in increasing the surface area (Figure 8-A), which in turn enhances the pre-mRNA splicing, as shown in Figure 8-B. This is because, the higher surface area, increases the probability of splicing particles to diffuse into the nuclear speckles resulting in increased localization. However, beyond ∼ 50 speckles the number of produced mRNA plateaus which could be due to the compensation of splicing particle localization by relatively smaller-sized nuclear speckles. Production of mRNA was maximized when there were between 20 and 50 speckles, which coincides directly with the experimentally determined values (18). In addition, the size of the nuclear speckles corresponding to the maximum mRNA production falls between 1.4 to 1 *μ*m, which is also compatible to the known nuclear speckles diameters of one to a few microns (18). Therefore, our results suggest that it is plausible that the cells optimize the design parameters of speckles (the number and size) to maximize the mRNA production.

**Fig. 8.**
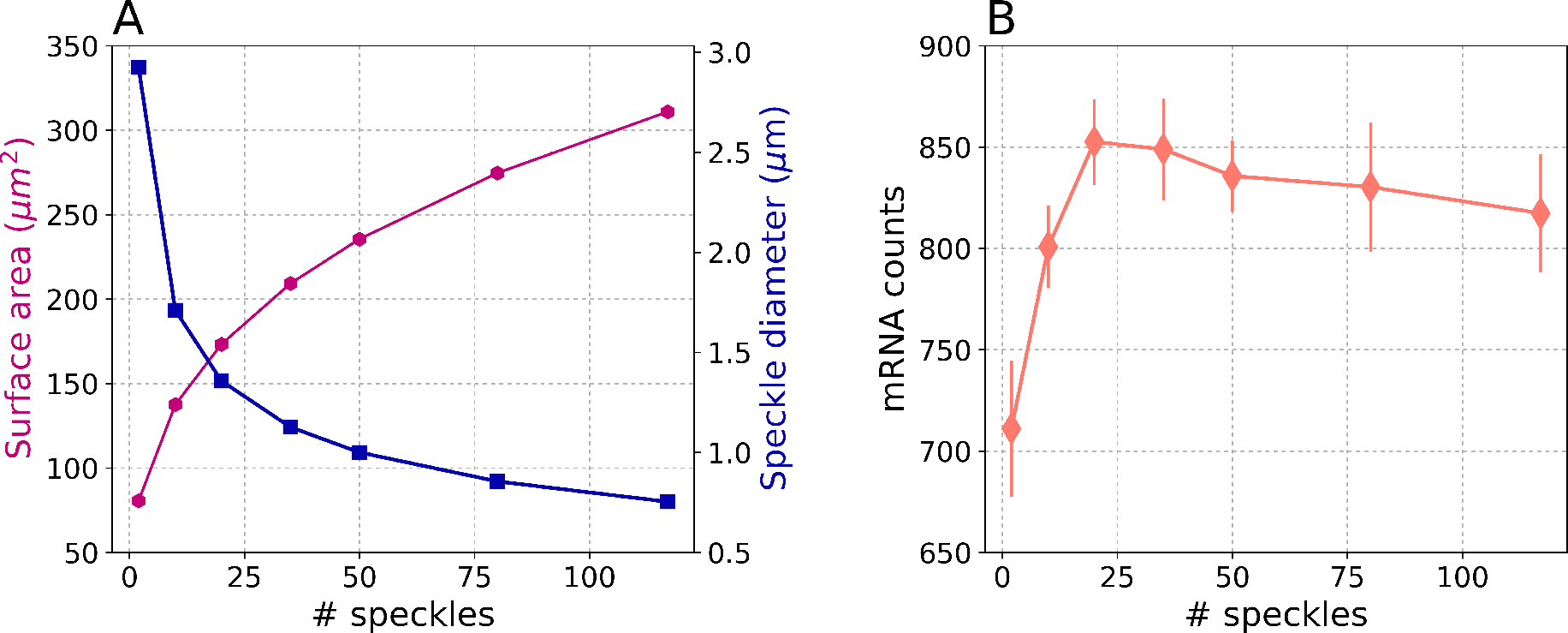
A) Increasing the number of nuclear speckles, results in an increase of the surface area (magenta curve) and decrease of the speckles diameter (blue curve); B) mRNA production increases as the number of speckles increases till about 50 speckles beyond which the production plateaus. Error bars represent the standard deviations. For each condition, 20 simulation replicates were performed.

### Gene distribution around speckles affect transcript splicing and mRNA production

It is known that genes are organized nonrandomly around nuclear speckles (31, 32). In a recent study, Chen et al. investigated the organization of whole genome using TSA-seq method (33). They showed that the most highly expressed genes are located between ≈ 0.05 and *μ*m from the periphery of a speckle. They also speculated that the genome movement of several hundreds of nano meter from nuclear periphery towards speckles could have functional significance. To test their hypothesis, we investigated the effects of active genes distribution around speckles’ periphery. We varied the gene distance from 0.054 to 2 *μ*m and observed the effects on the number of spliced mRNA transcripts found in cytoplasm. As Figure 9 demonstrates, increasing the distance of the genes to speckle periphery from ≈ 0.05 to 0.2 *μ*m sharply decreases the mRNA counts by a factor of 2, with no further significant decrease at larger distances. Thus, the effect can be even more pronounced over a short distance range than they were able to resolve. Considering the fact that our speckle model does not involve any active recruiting of the pre-mRNA transcripts, nor do our speckles move toward an active transcription site, the observed effect is mainly due to the diffusion of the transcripts in the nucleus before they become associated with the speckles. Our model predicts that the gene distribution around speckles has an effect on mRNA splicing. It is plausible that this effect might be regulated by speckle movement towards transcriptionally active genes, consistent with the fluid nature of these nuclear bodies.

**Fig. 9.**
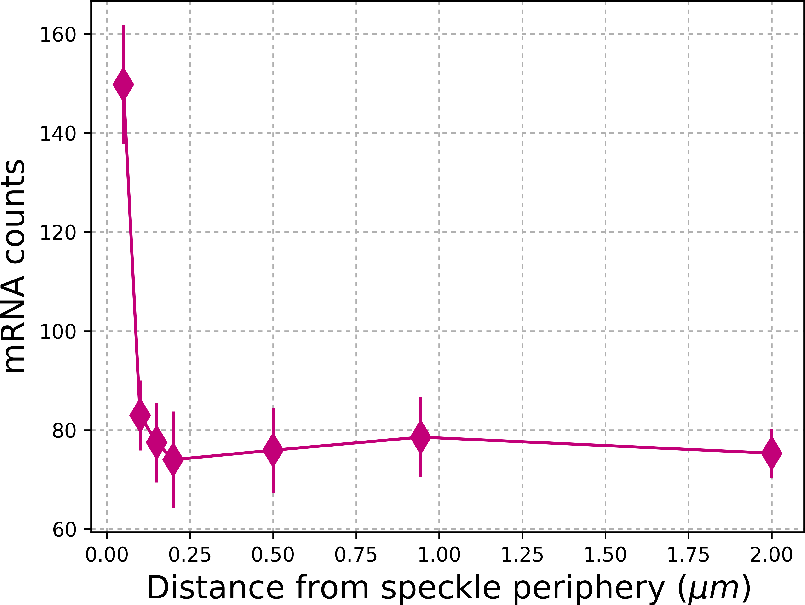
Our model predicts that the mRNA production decreases by a factor of two, above 0.054 *μ*m. Error bars represent the standard deviations. For each condition, 20 simulation replicates were performed.

## Conclusions

Spatial organization is one of the key features of eukaryotic cells that brings order to complex biochemical reactions. We studied two aspects of this organization both in connection with RNA splicing (the process of removal of non-coding regions from pre-mRNA transcripts). We investigate, firstly, the effects of cellular organelles on assembly of the building blocks of splicing; and secondly, mRNA production at nuclear speckles which are liquid organelles (droplets) that have higher concentrations of reactants involved in splicing. We constructed a spatially-resolved mammalian whole-cell (HeLa cell) model from a variety of experimental data. Using this model, we simulate the splicing event with a kinetic model we developed, as stochastic reaction-diffusion processes. We show that number of nuclear pore complexes—assemblies of proteins embedded in the nuclear envelope that control the traffic between nucleus and cytoplasm—affects the splicing particles formation rate for different nuclear sizes. The phase-separated nature of speckles enhances the mRNA production and reduces its associated noise. We suggest a rationale for the number and size of speckles based on optimal resulting mRNA. By demonstrating the effects of the genes distribution around our non-fluid speckles on mRNA counts, we propose that the movement of nuclear speckles toward active chromatin regions could be regulated by cells to control the transcripts production. Overall, our reaction-diffusion model of spliceosome assembly and function in a realistic mammalian cell environment allowed us to meaningfully connect the cellular geometry to the underlying biological processes. At the same time, we expect the presented whole-cell model to provide a versatile platform for studying processes beyond mRNA splicing.

## Methods

### Construction of a representative HeLa whole-cell model

Spliceosomal assembly and activity consists of multi-compartmental, reaction-diffusion processes, necessitating a spatial representation of the cellular geometry. HeLa cells are an optimal model system as they have been the subject of extensive investigations exploring cell geometry and cellular composition. Additionally, data from individual measurements of specific components (e.g., size, morphology, relative mass fraction) were used to inform the construction of a representative cell model (11, 14–20, 23–26). A constructive solid geometry (CSG) approach, wherein basic geometric objects are combined programmatically via set operations (e.g. unions, differences, intersections), was used to build the HeLa cell. Since Lattice Microbes software (v 2.3) (21, 22) requires that each location within the space be defined as a single site-type, the various CSG objects were stenciled onto the simulation lattice in “depth order” (also called the “Painter’s algorithm”). Overall, our model consists of 11 different site-types including: 1) extracellular space, 2) plasma membrane, 3) cytoplasm, 4) nuclear membrane, 5) nucleoplasm, 6) Cajal bodies, 7) nuclear speckles, 8) nuclear pore complexes, 9) mitochondria, 10) Golgi apparatus and 11) endoplasmic reticulum (ER). The overall simulation volume was constructed as a cubic box with 18.432 *μ*m side-length. The space was discretized into a cubic lattice of points spaced 64 nm apart. HeLa cell volumes have been measured at 2600, 3000 and 4400–5000 *μm*^3^ (24, 43, 44); we chose the mid-size cell as a template for our model. HeLa cells that are grown in suspension appear spherically-shaped, so we chose to design the overall cell architecture as a sphere with radius 8.9 *μ*m. Nuclei have measured volumes of, 220 and 374 *μm*^3^ (25, 26). Refs. (24) and (15) suggested nuclear volumes corresponding to 10% and 21.1% of the total cell volume. As we wanted to test the importance of nuclei size on splicing, multiple nuclear radii were investigated, including 3.74, 4.15, 4.67 and 5.29 *μ*m corresponding to all the above-motioned volumes and volume-fractions. Plasma and nuclear membranes were implemented as a thin sheet of lattice points (128nm thick) separating the extracellular space, the cytoplasm and the nucleus. The Golgi apparatus was constructed as an intersection of a cone with several spherical shells of various radii placed successively from the edge of the nucleus into the cytoplasm. The apex of the cone was centered in the cell with the based positioned deep in the cytoplasm. In this way, the Golgi roughly approximates what is seen in experiments. Nuclear speckles and Cajal bodies were modeled as spheres placed within the nucleus. Mitochondria were modeled as randomly oriented spherocylinders placed within the cytoplasm. Nuclear pore complexes were embedded in the nuclear membrane. NPCs were constructed as a set of spheres of radii approximately equal to that of the experimentally-measured NPC. Sizes for these organelles can be found in Table 1. Total counts for these organelles were based on either direct experimental quantification or based on relative volume fraction measured for the overall cell. The ER was also constructed in a randomized fashion with the details and construction algorithm presented in Supplementary Information. The endosomes, lysosomes, actin-cytoskeleton, peroxisomes in the cytoplasm; nucleolus and chromatin have not been included into the present version of the model. According to Ref. (15), each of the cytoplasmic organelles contribute less than 1% of to the total cell composition, and therefore, were not modelled. The nuclear components were chosen mainly among those that play a role in RNA splicing processes. A representative HeLa cell geometry resulting from this procedure is shown in Figure 1. The associated code for setting up a HeLa cell model for Lattice Microbes software (21, 22) is available upon request.

In addition to structural features, the abundance of the proteins participating in the processes we studied were derived from proteomics data of HeLa cells (19). The number of active snRNA genes have been determined to be 30 (45). These abundances were used as the initial condition for the simulations. Particles are randomly distributed throughout their parent region with locations sampled from a uniform distribution; for instance, genes are distributed throughout the nucleus. For each separate simulation replicate, a different initial particle placement was used.

### Kinetic models

**Table 2.** Reactions describing U1snRNP and U2snRNP splicing particles formation together with their associated rates. Abbreviations are: DNA(D), Gemin 5(*G*^5^), five already-assembled Sm proteins (*Sm*^5^), the remaining Sm proteins (*Sm*^2^), diffusion-limited (D.L.), model assumption (M), nucleus (N), NPC (P), Cajal bodies (J) and cytoplasm (C)

| Reaction | Rate | Units | Reference | Compartment |
| --- | --- | --- | --- | --- |
| **In Nucleus** |  |  |  |  |
| $D \rightarrow D + U1snRNA_{nuc}$ | 0.285 | $s^{-1}$ | (46) | N |
| $D \rightarrow D + U2snRNA_{nuc}$ | 0.224 | $s^{-1}$ | (46) | N |
| **Nucleus to Cytoplasm** |  |  |  |  |
| $U1(2)snRNA_{nuc} \rightarrow U1(2)snRNA_{cyt}$ | $2 \times 10^4$ | $s^{-1}$ | M | P |
| **Cytoplasmic Assembly** |  |  |  |  |
| $U1(2)snRNA_{cyt} + G^5 \rightarrow U1(2)snRNA \cdot G^5$ | $1.02 \times 10^8$ | $M^{-1}s^{-1}$ | D.L. | C |
| $U1(2)snRNA \cdot G^5 \rightarrow U1(2)snRNA_{cyt} + G^5$ | 3.05 | $s^{-1}$ | (47) | C |
| $U1snRNA \cdot G^5 + Sm^5 \rightarrow U1snRNA \cdot Sm^5 + G^5$ | $5.9 \times 10^7$ | $M^{-1}s^{-1}$ | D.L. | C |
| $U2snRNA \cdot G^5 + Sm^5 \rightarrow U2snRNA \cdot Sm^5 + G^5$ | $1.18 \times 10^7$ | $M^{-1}s^{-1}$ | D.L. | C |
| $U1(2)snRNA \cdot Sm^5 + Sm^2 \rightarrow U1(2)snRNA \cdot Sm^7$ | $1.39 \times 10^8$ | $M^{-1}s^{-1}$ | D.L. | C |
| $U1(2)snRNA \cdot Sm^7 \rightarrow U1(2)snRNA \cdot Sm^5 + Sm^2$ | 2.78 | $s^{-1}$ | (47) | C |
| **Cytoplasm to Nucleus** |  |  |  |  |
| $U1(2)snRNA \cdot Sm^7 \rightarrow U1(2)snRNA \cdot Sm^7_{nuc}$ | $2 \times 10^4$ | $s^{-1}$ | M | P |
| **Nuclear Maturation** |  |  |  |  |
| $U1snRNA \cdot Sm^7_{nuc} + U1_{prot} \rightarrow U1snRNP$ | $1.22 \times 10^7$ | $M^{-1}s^{-1}$ | (48) | J-N |
| $U1snRNP \rightarrow U1snRNA \cdot Sm^7_{nuc} + U1_{prot}$ | $4.8 \times 10^{-4}$ | $s^{-1}$ | (48) | J-N |
| $U2snRNA \cdot Sm^7_{nuc} + U2_{prot} \rightarrow U2snRNP$ | $0.24 \times 10^7$ | $M^{-1}s^{-1}$ | (48) | J-N |
| $U2snRNP \rightarrow U2snRNA \cdot Sm^7_{nuc} + U2_{prot}$ | $4.8 \times 10^{-4}$ | $s^{-1}$ | (48) | J-N |

**Table 3.** Spliceosome assembly and splicing reaction. Abbreviations are: DNA (D), U1snRNP (U1), U2snRNP (U2), U4/U6 U5snRNP (tri․ U), model assumption (M), diffusion-limited (D.L.), nuclear speckles (S) and nucleus (N)

| Reaction | Rate | Units | Reference | Compartment |
| --- | --- | --- | --- | --- |
| $D \rightarrow D + pre - mRNA$ | $4.7 \times 10^{-3}$ | $s^{-1}$ | (7) | S-N |
| $U1 + pre - mRNA \rightarrow complexE$ | $4.66 \times 10^7$ | $M^{-1}s^{-1}$ | D.L. (11) | S-N |
| $complexE \rightarrow U1 + pre - mRNA$ | 1.57 | $s^{-1}$ | (49) | S-N |
| $U2 + pre - mRNA \rightarrow complexE^*$ | $1.4 \times 10^7$ | $M^{-1}s^{-1}$ | D.L. (11) | N |
| $U2 + pre - mRNA \rightarrow complexE^*$ | $0.93 \times 10^7$ | $M^{-1}s^{-1}$ | D.L. (11) | S |
| $complexE^* \rightarrow U2 + pre - mRNA$ | 1.57 | $s^{-1}$ | (49) | S-N |
| $complexE + U2 \rightarrow complexA$ | $8.8 \times 10^7$ | $M^{-1}s^{-1}$ | D.L. (11) | S-N |
| $complexA \rightarrow complexE + U2$ | 0.062 | $s^{-1}$ | (49) | S-N |
| $complexE^* + U1 \rightarrow complexA$ | $8.7 \times 10^7$ | $M^{-1}s^{-1}$ | D.L. (11) | S-N |
| $complexA \rightarrow complexE^* + U1$ | 1.57 | $s^{-1}$ | (49) | S-N |
| $complexA + tri \cdot U \rightarrow complexB$ | $4.66 \times 10^7$ | $M^{-1}s^{-1}$ | D.L. (11) | S-N |
| $complexB \rightarrow complexA + tri \cdot U$ | 1.55 | $s^{-1}$ | (49) | S-N |
| $complexB \rightarrow complexB^* + U1$ | $6 \times 10^4$ | $M^{-1}s^{-1}$ | M | S-N |
| $complexB^* \rightarrow mRNA + tri \cdot U + U2$ | 0.067 | $s^{-1}$ | (50) | S-N |

### Implementation of nuclear speckles

Nuclear speckles were modelled as spherical regions in the nucleus with radii of 0.35 *μ*m. Previously, the splicing particles localization in speckles has been implemented by assuming a higher affinity for splicing particles to bind to unknown binding partners in speckles with respect to binding to pre-mRNA transcripts in the nucleoplasm (41). We imposed an imbalance on the transition probabilities for the splicing particles and pre-mRNA transcripts to move from the nuclear speckles to the nucleoplasm, and vice-versa. This approach, as shown in the Results section, will reproduce experimentally observed concentration ratio of splicing particles between the speckles and the nucleoplasm (41), additionally, the presence of dummy particles in the speckles are not required. Specifically, the probability of the splicing particles to move from the nucleoplasm regions to the speckles (*P*_*n*_) was higher than the reverse direction (*P*_*s*_). To examine the effect of the bias, the *P*_*n*_/*P*_*s*_ values were varied. With increasing *P*_*n*_/*P*_*s*_ values, more particles accumulate in the speckles and the nucleus becomes more diluted. Figure 10 shows the localization of splicing particles in speckles upon application of this bias in our model.

**Fig. 10.**
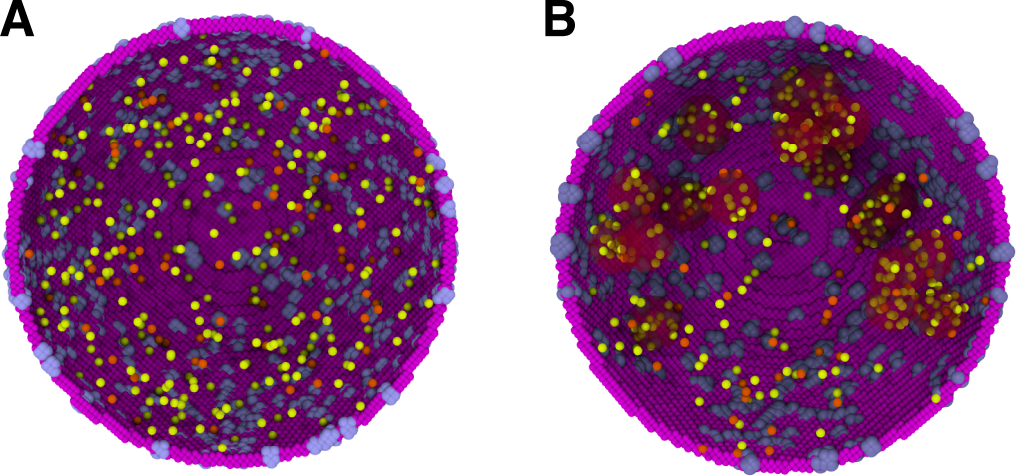
Formation of speckles in our simulations: A) splicing particles (U1 and U2 colored in yellow and orange, respectively) diffusing freely in the nucleus without speckle, B) Introduction of an imbalance on transition probabilities of splicing particles from the nucleus to speckles results in the localization of the splicing particles in the speckles shown as red-shaded regions.

## Acknowledgements

This work was supported by the National Science Foundation [grant MCB-1244570] to Z.G. and Z.L-S, Graduate Fellowship Program [grant DGE-1144245] to J.R.P. and [grant MCB 1803786] to M.G‥ Z.L-S held the William and Janet Lycan Chair in Chemistry and M.G. held the James R. Eiszner Chair while this work was carried out. Supercomputer time was provided by XStream-XSEDE [grant TG-MCA03S027]. Z.G. thanks Prof. Andrew Belmont and Dr. Jospeph Dopio (MCB, UIUC) for many spontaneous and stimulating conversations.

Z.G. thanks Prof. Andrew Belmont, Prof. Prashant Jain (Chemistry, UIUC), Prof. K. Prasanth (MCB, UIUC), Dr. Marian Breuer (Chemistry, UIUC) and Dr. Pankaj Chaturvedi (MCB, UIUC) for critical reading of the manuscript and helpful comments. Z.G. especially thanks Mike Hallock (School of Chemical Sciences, UIUC) for support with Lattice Microbes code.

Z.G, J.R.P., M.G, and Z.L-S. designed research; Z.G. and J.R.P. performed research; Z.G. analyzed data; and Z.G, J.R.P., M.G, and Z.L-S. wrote the paper. All authors read and approved the manuscript.

